# CRISPR/Cas9-mediated generation of biallelic G0 anemonefish (*Amphiprion ocellaris*) mutants

**DOI:** 10.1101/2020.10.07.330746

**Authors:** Laurie J. Mitchell, Valerio Tettamanti, Justin N. Marshall, Karen L. Cheney, Fabio Cortesi

## Abstract

Genomic manipulation is a useful approach for elucidating the molecular pathways underlying aspects of development, physiology, and behaviour. However, a lack of gene-editing tools appropriated for use in reef fishes has meant the genetic underpinnings for many of their unique traits remain to be investigated. One iconic group of reef fishes ideal for applying this technique are anemonefishes (Amphiprioninae) as they are widely studied for their symbiosis with anemones, sequential hermaphroditism, complex social hierarchies, skin pattern development, and vision, and are raised relatively easily in aquaria. In this study, we developed a gene-editing protocol for applying the CRISPR/Cas9 system in the false clown anemonefish, *Amphiprion ocellaris*. Microinjection of eggs at the one-cell stage was used to demonstrate the successful use of our CRISPR/Cas9 approach at two separate target sites: the rhodopsin-like 2B opsin encoding gene (*RH2B*) involved in vision, and Tyrosinase-producing gene (*tyr*) involved in the production of melanin. Analysis of the sequenced target gene regions in *A. ocellaris* embryos showed that uptake was as high as 50% of injected eggs. Further analysis of the subcloned mutant gene sequences revealed that our approach had a 75% to 100% efficiency in producing biallelic mutations in G0 *A. ocellaris* embryos. Moreover, we clearly show a loss-of-function in *tyr* mutant embryos which exhibited typical hypomelanistic phenotypes. This protocol is intended as a useful resource for future experimental studies that aim to elucidate gene function in anemonefishes and reef fishes in general.

## INTRODUCTION

Targeted genome modification (i.e. reverse genetics) is an elegant approach for directly attributing genotype with phenotype but has been limited in non-model organisms owing to a lack of high-quality assembled genomes, affordable technology and species-specific protocols. Modern gene-editing tools such as the clustered-regularly-interspaced-short-palindromic-repeat (CRISPR/Cas9) system enables precise targeted gene-editing, where a synthetic guide RNA (sgRNA) directs the cutting activity of Cas9 protein to produce a double strand break at a genetic location of interest. Subsequent error prone DNA repair by non-homologous end joining (NHEJ) often leaves insertions and/or deletions (indels), which may induce a frameshift and potential loss of gene function (Hsu, Lander & Zang, 2014). The injection of sgRNA fused with Cas9 protein has proven to be an effective tool for precise genome editing at target gene sequences in the cell lines of numerous species including many teleost fishes such as zebrafish (*Danio rerio*) (for a review, see Li et al. 2016), Nile tilapia (*Oreochromis niloticus*) (Li et al. 2014; Zhang et al. 2014), medaka (*Oryzias latipes*) (Ansai & Kinoshita, 2014), Atlantic salmon (*Salmo salar*) (Edvardsen et al. 2014), killifish (spp.) (Aluru et al. 2015; Harel et al. 2015), pufferfish (*Takifugu rubribes*) (Kato-Unoki et al. 2018), and red sea bream (*Pagrus major*) (Kishimoto et al. 2018). However, the CRISPR/Cas9 system has yet to be applied to coral reef fishes, a highly diverse assemblage of species with a unique life-history (Cowen & Sponaugle, 1997) and multitude of biological adaptations (Peterson & Warner, 2002; Wainwright & Bellwood, 2002) suited for survival in their marine environment.

One such group of reef fishes suitable for gene-editing are anemonefishes (subfamily, Amphiprioninae), an iconic group that shelter in certain species of sea anemones (Fautin & Allen, 1997), and are sequential hermaphrodites (Fricke, 1983; Ochi, 1989) that live in strict social hierarchies determined by body size (Buston, 2003). The fascinating aspects of anemonefish biology has led to their use in multiple areas of research including for studying the physiological responses of reef fishes to the effects of climate change (Scott & Dixson, 2016; Beldade et al. 2017; Norin et al. 2018), the hormonal pathways that regulate sex change (Casas et al. 2016; Dodd et al. 2019) and parental behaviour (DeAngelis et al. 2017, 2018; Iwata & Suzuki, 2020), and the physiological adaptations for group-living (Buston, 2003; Buston & Cant, 2006). Moreover, anemonefishes are being used to understand the visual capabilities of fish (Stieb et al. 2019; Mitchell et al. 2020) and evolution of skin colour diversity (Maytin et al. 2018; Salis et al. 2018; Salis et al. 2019) in reef fishes. However, despite the wide-reaching applications of anemonefish research, the genetic basis for many of their traits remain to be empirically investigated. Consequently, anemonefish studies have been limited to correlative findings from comparative transcriptomics (Maytin et al. 2018; Salis et al. 2018, 2019) and/or indirect comparisons by using reverse genetics/testing genetic elements of interest in pre-established models such as zebrafish (e.g. Salis et al. 2019). Recently, the release of assembled genomes for multiple anemonefish species (Tan et al. 2018; Lehmann et al. 2019; Marcionetti et al. 2019) has made it feasible to apply the CRISPR/Cas9 system to conduct genome modification in anemonefishes.

Producing biallelic knockout animals within the first (G0) generation is often essential for the development of transgenic animal lines, particularly in species with long generation times, and requires a well-designed protocol for the efficient delivery of CRISPR/Cas9 constructs to completely knockout gene function and minimise the chance of chimerism/mosaicism. To achieve this, careful species-specific considerations must be made for sgRNA design, dose toxicity, construct delivery parameters (i.e. air pressure, needle dimensions), and egg/embryo-care both during microinjection (e.g. Kishimoto et al. 2019) and incubation. Inherent challenges specific to gene-editing anemonefishes and many other demersal spawning reef fishes include the injection and/or care of their substrate-attaching eggs (Roux et al. 2019) and pelagic larval stage (Leis & McCormick, 2002). Therefore, a protocol for performing CRISPR/Cas9-mediated genome editing in anemonefishes would be highly beneficial for diverse areas of research to directly test candidate genes facilitating sex change (Dodd et al. 2019), colour vision (Mitchell et al. 2020) and skin pattern development (Salis et al. 2019).

In this study, we describe a protocol for performing CRISPR-Cas9 in the false clown anemonefish, *Amphiprion ocellaris*, an ideal species for gene-editing due to the public availability of its long-read assembled genome (Tan et al. 2018), its relative ease of being cultured in captivity (Mazzoni et al. 2019), and being the most widely studied anemonefish species (for a review, see Roux et al. 2020). To demonstrate our protocol, we report on its efficacy in producing biallelic knockouts in G0 generation *A. ocellaris* injected with synthetic guide RNA and recombinant Cas9 protein that separately targeted two genes, including the rhodopsin-like 2B opsin gene (*RH2B*) encoding a mid-wavelength-sensitive visual pigment (Bowmaker, 2008), and the Tyrosinase encoding gene (*tyr*) involved in the initial step of melanin production (Cal et al. 2017). Moreover, analyses of sequencing results and skin (melanism) phenotype from embryos revealed in many individuals a complete loss of gene function. We hope this protocol provides a useful resource for future gene-editing experiments in anemonefishes and similar demersal spawning reef fishes.

## MATERIALS AND METHODS

### Care and culturing of *A. ocellaris*

Captive-bred pairs of *A. ocellaris* purchased from a local commercial breeder (Clownfish Haven, Brisbane Australia) were housed in recirculating aquaria at The Institute for Molecular Bioscience at The University of Queensland, Australia. Experiments were conducted in accordance with Animal Ethics Committee guidelines and governmental regulations (AEC approval no. QBI/304/16; Australian Government Department of Agriculture permit no.

2019/066; UQ Institutional Biosafety approval no. IBC/1085/QBI/2017). Individual aquaria (95 L) contained a single terracotta pot (27 cm diameter) that provided a shelter and egg-laying structure for brood-stock fish. Spawning usually occurred during the late-afternoon to evening (15:00-18:00), which was preceded by a fully protruded ovipositor and behaviours that included surface cleaning and ventral rubbing on pot surfaces. Eggs laid by the female were adhered to the pot and subsequently fertilised by the male. Eggs were incubated in an isolated tank (36 L) that contained heated (26°C) marine water (1.025 sg) dosed with methylene blue (0.5 mL, Aquasonic), and were kept aerated using a wooden air diffuser (Red Sea). Although this study did not analyse mutagenesis beyond the embryo stage (Fig. 1B), we have included a detailed guide on larval hatching and rearing in the supplementary materials.

**Figure. 1.**
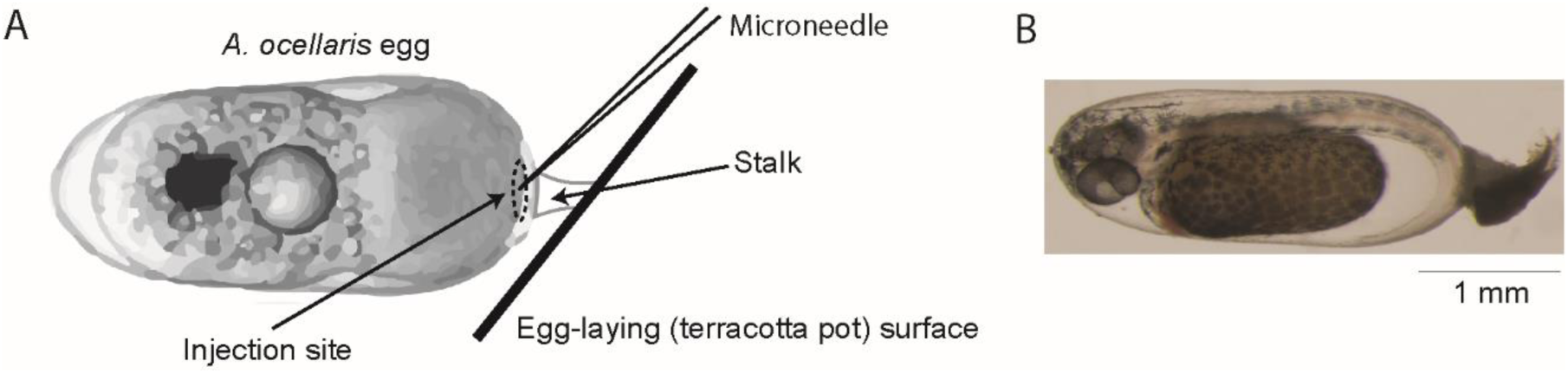
(A) Illustration depicting an *A. ocellaris* egg (less than 1 h post-fertilisation) adhered to substrate (terracotta pot) by a fibrous stalk. Circled at the substrate-facing side of the egg is the injection site at the animal pole where microinjections were aimed. (B) Wild-type *A. ocellaris* embryo at 64-88 hours post-fertilisation with formed eyes and melanin pigmentation.

### Design and in-vitro testing of sgRNAs

To trial the application of the CRISPR-Cas9 system in anemonefishes, we designed three and two sgRNAs that targeted *A. ocellaris RH2B* and *tyr* genes, respectively (Fig. 2A, B). The gene sequence for *A. ocellaris RH2B* was obtained from a previous study (Mitchell et al. 2020), and the same approach described by Mitchell et al. 2020 was used to identify the *tyr* gene sequence in the *A. ocellaris* genome (Tan et al. 2018). All gene sequences were viewed in Geneious (v.2019.2.3), and the “Find CRISPR Sites…” function was used to screen suitable sgRNA sequences with search parameters that included a target sequence length of 19-bp or 20-bp, an ‘NGG’ protospacer-adjacent-motif (PAM) site located on the 3’ end of the target sequence, and off-target scoring against the *A. ocellaris* genome (see supplementary material for a list of sgRNA sequences). All selected target sequences were screened to ensure no major off-target sites were present (≥90% specificity). Both the sgRNAs and purified Cas9 protein used in this study were purchased from Invitrogen (catalogue no. A35534, A36498). One forward-directed cutting sgRNA on the *RH2B* gene targeted a sequence on Exon 4 immediately upstream (18-bp) of the conserved chromophore binding site Lys296 (Palczewski, 2006), where a frameshift would prevent the formation of a functional visual pigment. To assess cutting activity at other *RH2B* sites, we selected two additional target sequences on Exon 5 (i.e. downstream of Lys296), that may allow future attempts to remove the entire binding site by co-injecting sgRNA. Two sgRNAs targeted sites on Exon 2 of the *tyr* gene, a location adequately upstream where reading frame shifts produced by indel mutations would more likely knockout gene function, while being far enough downstream to reduce the likelihood of alternative transcription start sites being utilised. The cutting activity of our sgRNAs with Cas9 were initially assessed *in-vitro* by incubating PCR amplicons of each targeted gene region with or without sgRNA and/or Cas9 and comparing fragment length via gel electrophoresis (see supplementary material for full details on PCR routine, reagent quantities and incubation parameters) (Fig. 2C, D).

**Figure. 2.**
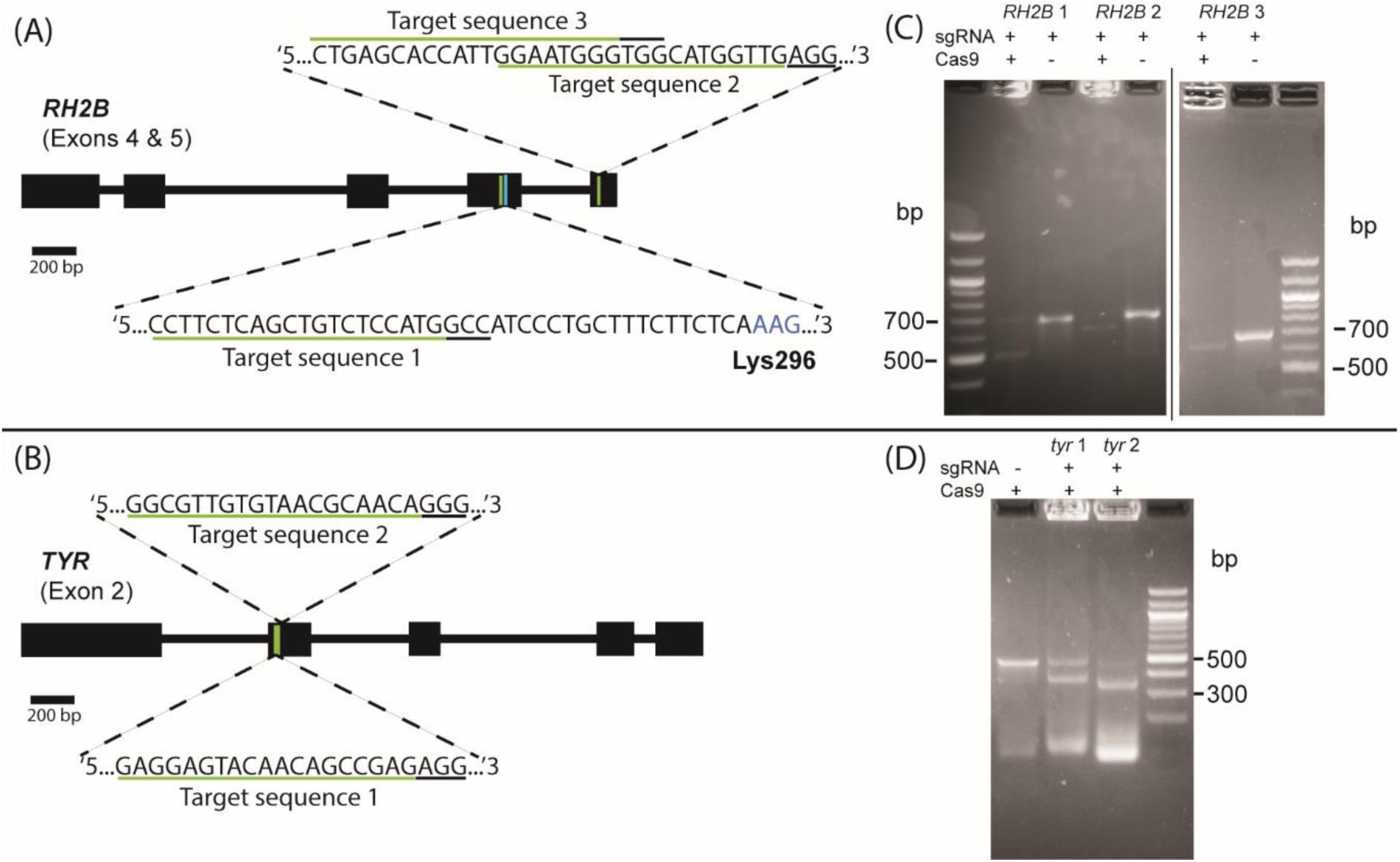
Sites and sequences targeted by sgRNA designed for the *RH2B* (A) and *tyr* (B) genes in *A. ocellaris*. Expanded regions show the target sequence (underlined in green) and ‘NGG’ PAM (underlined in black) for each sgRNA. For Exon 4 of *RH2B*, the Lys296 chromophore binding site (coloured blue) is also depicted down-stream of target sequence 1. Gel images to the right of each gene illustration depict DNA fragments size when amplicons that contained targeted (C) *RH2B* and *tyr* (D) gene regions were incubated (*in-vitro*) with or without Cas9 protein and sgRNA.

### Microinjection delivery of CRISPR-constructs

The clutches were collected 10-15 minutes post-fertilisation for CRISPR-construct delivery to ensure adequate fertilisation of eggs but before the first cell division had occurred (60 – 90 min post-fertilisation (Yasir & Qin, 2007)). Pots containing egg clutches were broken apart into multiple shards (∼2.0×4.0 cm) using a hammer and chisel. Post-delivery, the shards were mounted in a petri dish and partially submerged in Yamamoto’s ringer’s solution to prevent dehydration of eggs and osmotic stress associated with injection (Kinoshita et al. 2009; Kishimoto et al. 2019). Eggs were viewed under a dissection microscope (3.5x magnification) and microinjected directly into the animal pole at a 45° angle with a pulled borosilicate glass pipette (Harvard Apparatus: 1.0×0.58×100 mm) fitted on a pneumatic injector unit (Narishige IM-400) (Fig. 1A) and micromanipulator (Marzhauser MM3301R). Injector pressure settings were configured to deliver a 1 nL dose of a mixture per egg. The mixture contained sgRNA (200 ng/μL), Cas9 protein (500 ng/μL) and KCl (300 μm), was initially suspended by slowly pipetting up-and-down in a 10ul stock-solution containing 5.5 μL RNAse free H2O and incubated at 37°C for 10 minutes before being stored on ice, 20 – 30 min before injections started. Both the inclusion of KCl solution to aid in sgRNA/Cas9 mix solubility, and incubation step were adapted from Burger et al. (2016). 2 μL of the solution was then backloaded into a microneedle immediately before injection (see supplementary material for full details on microneedle dimensions and injector pressure settings). Injecting ceased when the chorion had become too thick to penetrate (∼40-50 minutes post-fertilisation). To assess the mortality attributed to toxicity of the injection dosage and damage induced loss, the survival rate of CRISPR-Cas9 injected eggs were compared to controls, including: *1*) eggs injected with a mixture containing no Cas9 (replaced with water), and *2*) non-injected eggs. To control for differences in individual user, we had multiple personnel perform injections across clutches. Survival rates for eggs were calculated as the proportion of live embryos at collection relative to the number of live eggs per treatment at <1 hour post-fertilisation (hpf).

### Genotype and phenotype analysis of mutant embryos

Treatment and control eggs were collected 64-88 hours post-fertilisation when live embryos with formed eyes were clearly visible (Fig. 1B). DNA was extracted from embryos using a DNeasy Blood & Tissue kit (Qiagen catalogue no. 69504), as per the manufacturer’s protocol. The concentration and purity of the extracted DNA was first tested via Nanodrop nucleic acid quantification and then PCR-amplified using primers flanking the targeted gene location (see supplementary material for gene-specific primer sequences). Sanger sequencing of PCR amplicons was outsourced to AGRF (https://www.agrf.org.au/) and positive mutants were detected by mapping their sequences against the respective gene in Geneious. Because all positive mutants were heterozygous, we identified the full range of mutations by subcloning the PCR products of four *RH2B* (clutch no. 3) and four *tyr* (clutch no. 7) mutants using the Invitrogen TOPO TA kit according to the manufactures protocol (Invitrogen catalogue no. K4575J10), and Sanger sequenced the extracted plasmids from 6-10 colonies per embryo (Fig. 3A, B). Mutants selected for cloning were also analysed via T7 endonuclease I-based (T7E1) heteroduplex assay according to the manufacturers protocol (EnGen® Mutation Detection Kit, NEB #E3321), and the length of digested and undigested fragments were visually compared by gel electrophoresis (Fig. 3C, D). Brightfield micrographs were taken (Nikon SMZ800N) of individual *tyr* mutant embryos and a wildtype embryo for comparison.

**Figure. 3.**
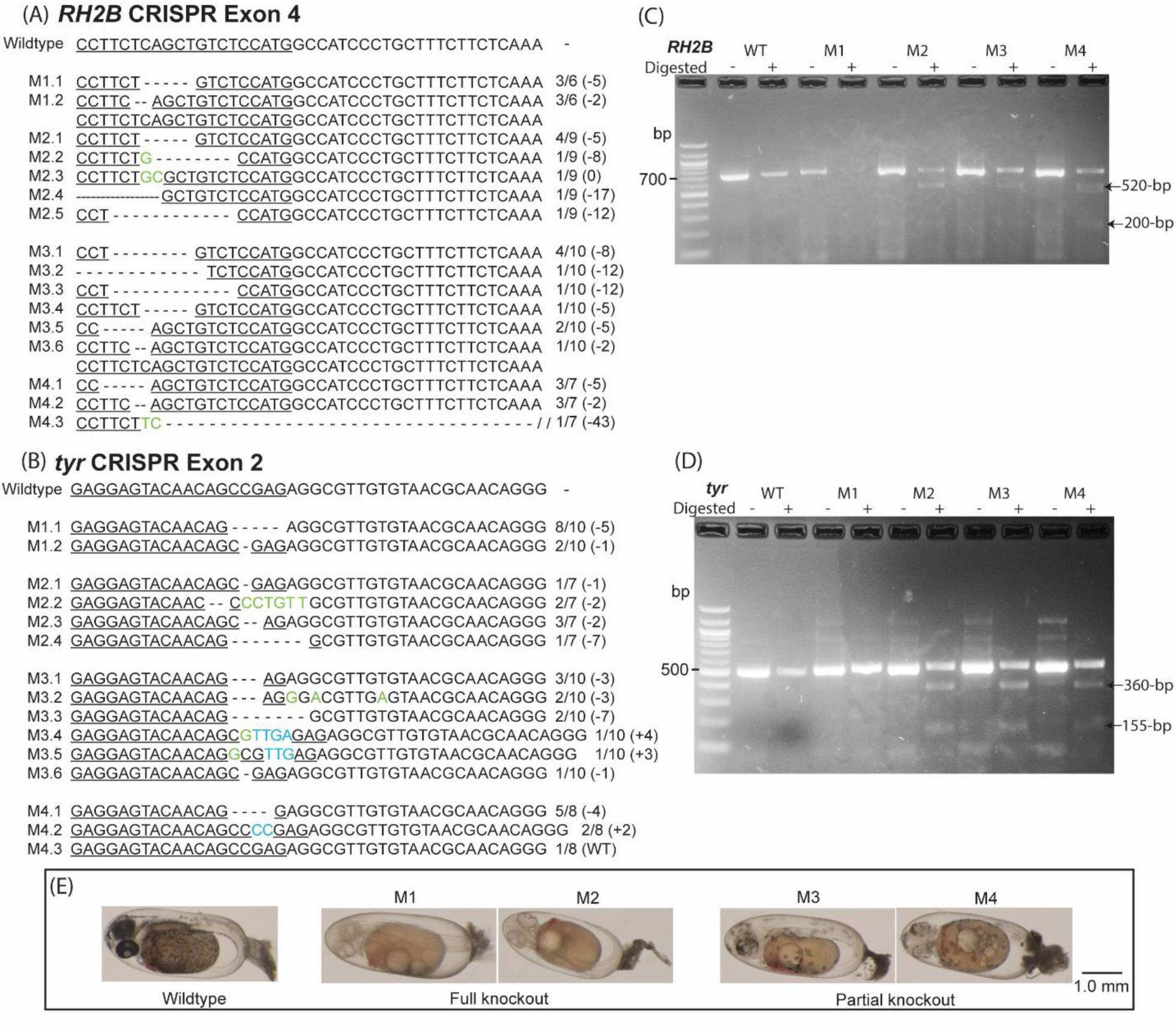
Subcloned sequences belonging to *A. ocellaris* embryos (clutch 3, *RH2B* 1; clutch 7, *tyr* 1) with mutations at targeted sequences (underlined) located on (A) Exon 4 of the *RH2B* opsin gene, and (B) Exon 2 of the *tyr* gene. Wildtype sequences are included as a reference. Detected mutations included deletions (dashes), substitutions (green), and insertions (blue). Sequence labels on the left-side indicate mutant and allele no., while numbers on the right-side indicate the detected frequency of each subcloned sequence in each embryo and the size of deletions (-) or insertions (+) is denoted in parantheses. Gel taken images of the T7E1 heteroduplex assay for (C) four *RH2B* and (D) four *tyr* mutants, with non-digested (homoduplex) and digested (heteroduplex) treatments, and wildtype (WT) treatments for reference. (E) Micrographs of *tyr* mutant *A. ocellaris* embryos exhibiting full knockout (*tyr*-M1 and 2) and partial knockout (*tyr*-M3 and 4) phenotypes, and a wildtype embryo for comparison.

## RESULTS AND DISCUSSION

### sgRNA *in-vitro* assay

An *in-vitro* assessment of sgRNA cutting activity was conducted to verify the integrity and viability of our sgRNA designs which targeted sites located on either *A. ocellaris RH2B* opsin gene (Fig. 2A) or *tyr* gene (Fig. 2B). All five selected sgRNAs exhibited positive cutting activity after incubation with amplicons that encompassed the targeted genes (Fig. 2C, D), although *tyr* 1 had a relatively low cutting efficiency, as evident by the near equally intense non-cleaved DNA band. Cutting activity indicated the sgRNA designs were suitable for *in-vivo* trials. No cutting activity was observed when amplicons were incubated without sgRNA (for *tyr*) or Cas9 (for *RH2B*).

### Survival and mutation rate

Overall, baseline (non-injected) clutch survival (mean ± sd: 62.2 ± 26.4%) was consistently higher than (sgRNA and Cas9) injected eggs (24.2 ± 8.6%), but inconsistently differed from sgRNA-only injected treatments, where it was lower in clutch 4 (Table 1). These observed differences in survival between the injected treatments and (non-injected) control eggs, indicates that physical trauma from the injection process was the major contributor to mortality observed in injected eggs. A reduction in needle tip-size (<15 μm) may help lower mortality; however, in our experience thinner needles exhibited excessive bending when attempting to penetrate the thick chorion of anemonefish eggs. Only needles with a relatively short-taper and broad tip (i.e. stubby profile) were usable for injections. Natural thickening of the chorion peaked at 30 – 40 minutes post-fertilisation (about 50 – 60 minutes preceding the first cell division) and prohibited further injecting.

**Table. 1.** Survival rates of injected and non-injected eggs at time of collection (64-88 hpf), number of genotyped embryos, and mutation rate per clutch and target sequence. “hpf” = hours post-fertilisation.

| Treatment/target sequence | Clutch 1 | Clutch 2 | Clutch 3 | Clutch 4 | Clutch 5 | Clutch 6 | Clutch 7 | Clutch 8 |
| --- | --- | --- | --- | --- | --- | --- | --- | --- |
|  | *RH2B* 1 | *RH2B* 1 | *RH2B* 1 | *RH2B* 2 | *RH2B* 2 | *RH2B* 3 | *tyr* 1 | *tyr* 2 |
| Injected survival 64-88 hpf | 11/50 (22.0%) | 14/48 (29.2%) | 13/48 (27.1%) | 7/56 (12.5%) | 17/50 (34.0%) | 16/45 (35.6%) | 14/86 (16.3%) | 9/47 (19.1%) |
| No. of genotyped (out of injected) embryos | 11/11 | 14/14 | 13/13 | 7/7 | 17/17 | 9/16 | 14/14 | 8/9 |
| Positive mutants | 1/11 (9.1%) | 3/14 (21.4%) | 4/13 (30.8%) | 1/7 (14.3%) | 8/17 (47.1%) | 3/9 (33.3%) | 7/14 (50.0%) | 2/8 (25.0%) |
| Non-injected survival rate 64-88 hpf | 25/59 (42.4%) | 40/50 (80%) | 25/42 (59.5%) | 16/52 (30.8%) | 44/47 (93.6%) | 51/61 (83.6%) | 37/45 (82.2%) | 12/40 (30.0%) |
| Injected (sgRNA-only) survival 64-88 hpf | - | - | - | 4/10 (40.0%) | 5/12 (41.7%) | - | - | - |

Examination of the targeted gene sequences of injected embryos showed highly variable mutation rates (Table 1) that ranged from 9.1% to 47.1% for RH2B (n=6 clutches), and 25% to 50% for tyr (n=2 clutches). To achieve higher mutation rates, we suggest a couple ways to improve the accuracy of injecting the animal pole by enhancing its visibility including *1*) the use of neutral coloured pots (e.g. white, grey) to contrast against the orange of anemonefish eggs, and *2*) delay injecting till the formation and visible swelling of the blastodisc (∼60 minutes post-fertilisation) that precedes the first cell division, although this may not be feasible with thickening of the chorion. Alternatively, the substitution of Cas9 protein with Cas9 mRNA may circumvent the need for direct delivery into the nucleus and permit injection elsewhere (e.g. in the yolk). Although Cas9 protein has been associated with a higher efficiency of mutagenesis than Cas9 mRNA (Kotani et al. 2015), the relatively long-lived (∼90 minutes) single cell stage of the *A. ocellaris* zygote (Yasir & Qin, 2007) would likely permit adequate time for migration into the nucleus and transcription/translation processes. The incorporation of nuclear-localisation-signal-fused Cas9 mRNA could also help compensate for differences in uptake efficiency (Hu et al. 2018).

### Genotype analysis of mutants

Analysis of the subcloned sequences of RH2B (clutch 3, RH2B 1) and tyr (clutch 7, tyr 1) mutant *A. ocellaris* embryos, revealed that our approach was successful in producing biallelic mutations in seven out of the eight embryos; only one tyr mutant retained a wildtype allele (Fig. 3A,B). This high efficiency (75% to 100%) in inducing biallelic mutations in G0 A. ocellaris fulfils a requirement for rapid reverse-genetic experiments that circumvents the need for backcrossing to establish a homozygous-line; often not feasible, particularly in anemonefish that take 12 – 18 months to reach sexual maturity (Madhu, Madhu & Retheesh, 2012).

Both *RH2B* (Fig. 3A) and *tyr* (Fig. 3B) mutant embryos had between two to six distinct mutations. This high number of mutations per embryo suggests Cas9 cutting activity persisted beyond the first cell division, an indication of a high dosage of sgRNA and Cas9 that could potentially be reduced if required. A total of 10 distinct mutations each were found in *RH2B* mutants (Fig. 3A) and in *tyr* mutants (Fig. 3 B), with most being in the form of deletions that ranged in length between 2 – 43bp and 1 – 7bp, respectively. Most mutations were situated (4 – 14bp) upstream (‘5) of their respective PAM sequence, a proximity and location near what is typically reported for Cas9 cutting activity (Jinek et al. 2012) (Fig. 3A, B). Exceptions included mutations in *tyr*-M2 and *tyr*-M3 with −7bp alleles starting at the PAM, and *RH2B*-M4 where a 43-bp deletion spanned regions both up- and down-stream of the PAM. The most frequent mutations found in multiple *RH2B* mutants included a 5bp deletion (10bp upstream of PAM) and a 2bp deletion (14bp upstream of PAM) (Fig. 3A), while the most common mutations across *tyr* mutants were a 1bp deletion (4bp upstream of PAM) and a 7bp deletion (starting at PAM).

A secondary analysis of mutant amplicons by T7E1 heteroduplex assay (Fig. 3C, D) exhibited digested (heteroduplex) DNA fragments for *RH2B* (520bp, 200bp; Fig. 3C) and *tyr* mutants (360bp, 155bp; Fig. 3D) that closely matched their amplicon lengths of 719bp and 512bp, respectively. Although there were no obvious digested fragments for *RH2B*-M1 (Fig. 3C), the faint non-digested (homoduplex) banding (∼700bp) suggests this was due to low nucleic acid input rather than lack of heteroduplex formation.

### Phenotype analysis of mutants

CRISPR/Cas9 knockout of *A. ocellaris tyr* produced seven embryos that exhibited varying degrees of hypomelanism (Fig. 3 E), a phenotype attributed to the disruption of the enzymatic conversion of tyrosine into melanin, and is similarly observed in *tyr* knockout zebrafish embryos and larvae (Ota & Kawahara, 2013; Jao, Wente & Chen, 2013). In comparison, wildtype *A. ocellaris* embryos consistently had heavily pigmented skin and eyes. A complete lack of melanin was observed in two (*tyr*-M1 and *tyr*-M2) out of the 14 injected embryos (Fig. 3E). Analysis of their subcloned sequences revealed both had biallelic mutations, all of which are likely to induce frameshifts that render TYR non-functional (Fig. 3B). Whereas partial depigmentation or a mosaic appearance was found in five out of the 14 embryos (e.g. *tyr*-M3 and *tyr*-M4; Fig. 3 E), most likely as a result of an incomplete knockout of TYR activity caused by in-frame mutations (*tyr*-M3.1, 3.2, 3.5), or heterozygosity (*tyr*-M4.3) from monoallelic cutting activity. The nature of this skin pigmentation phenotype has been shown in zebrafish to be sgRNA/Cas9 dose-dependent (Jao, Wente & Chen, 2013); however, in our case the nature of the mutation (i.e. in-frame or out-of-frame) was also a major determinant of phenotype. The low cutting efficiency of our sgRNA (*tyr* 1), as observed in the in-vitro cutting assay (Fig. 2D) may have also contributed to the more frequently observed incomplete knockout of TYR, by producing more monoallelic mutations in eggs.

Because there were no discernible phenotype(s) in *RH2B* mutant embryos, we are left to speculate on a loss of gene function based on the nature of the mutations (Fig. 3A). Two of the four subcloned *RH2B* mutants (*RH2B*-M1 and M4) possessed a full complement of mutant alleles that exhibited frameshifts and/or an extensive deletion encompassing the coding region (*RH2B*-M4.3). Examination of the translated (frameshifted) sequences confirmed the presence of missense mutations that disrupted the chromophore binding site (Lys296), and downstream premature stop codons may have precluded visual pigment formation (see Supplementary Figure 1). Thus, it is likely these two embryos had a total knockout of *RH2B* gene function. Behavioural experimentation will be necessary to demonstrate a functional loss of visual opsin in mutant anemonefish larvae/adults, as has been demonstrated in opsin knockout strains of medaka that exhibit impaired spectral sensitivity in optomotor tests (Homma et al. 2017) and/or altered social behaviour (Kamijo, Kawamura & Fukamachi, 2018; Kanazawa et al. 2020). Similarly, the loss of TYR could also be assessed for its impact on colour sensitivity, as has been reported in zebrafish (Park et al. 2016).

## Conclusion

Here we present the first use of the CRISPR/Cas9 system in a reef fish. Targeting the coding regions of the *RH2B* opsin and *tyr* genes successfully induced indel mutations in up to 50% of *A. ocellaris* embryos. Moreover, the analysis of subcloned sequences showed our gene-editing approach was able to produce biallelic mutations with an extremely high efficiency of ∼90%, causing complete loss-of-function mutations in a substantial proportion of G0 mutants. This opens the door to conducting gene-editing experiments in *A. ocellaris* to study the genetic basis for various anemonefish traits including sex change, skin pattern formation, parental behaviour, and vision. It also paves the way for similar approaches in other reef fish species. Our proven application of this technology to produce knock outs greatly facilitates the use of CRISPR/Cas9 for a variety of other genetic applications including making precise (knock-in) gene insertions in anemonefish.

## Author contributions

L. J. M., N. J. M, K. L. C and F. C. conceived the study. L. J. M., V. T., and F. C. designed guide RNAs, performed microinjections, and carried out the daily care of eggs. L. J. M ran the in-vitro cutting assay and T7E1 endonuclease assay. L. J. M. and V. T. performed subcloning and analysis of subcloned sequences. L. J. M wrote the initial manuscript, and all authors contributed to the final version of the manuscript.

## Acknowledgements

We thank Assoc. Prof. Justin Rhodes (University of Illinois, USA) for his assistance and generosity during an initial pilot study. We also thank the University of Queensland Biological Resources Aquatics Team, particularly Gillian Lawrence and Gerard Pattison for their support in maintaining marine aquaria and sourcing injection equipment.

## Funding

This research was funded by an Australian Research Council Discovery Project (DP18012363) awarded to N.J.M. and F.C. K.L.C was furthermore supported by an ARC Future Fellowship (FT190100313) and F.C. was supported by an ARC DECRA (DE200100620) and a University of Queensland Development Fellowship.

## Conflict of interest statement

The authors declare no conflicts of interest.

## Notes

### Competing Interest Statement

The authors have declared no competing interest.

### Summary of Updates

Included missing reference in methods (Burger et al. 2016).

