## supplementary for "CRISPR/Cas9-mediated generation of biallelic G0 anemonefish (*Amphiprion ocellaris*) mutants"

**Supplementary Material and Methods**

### 1. *A. ocellaris* egg hatching and larval rearing

Equipment required:

- 2x small aquaria (<50 L)
- Air pump, airline tubing and wooden air diffuser
- Black plastic wrap or plastic sheet material
- Paper or fabric towel
- Established rotifer culture
- Algal paste
- *Artemia* cysts and hatchery

For 0 to 6 days post-fertilisation (dpf), the eggs are kept in isolated aquaria (36 L volume) containing water with parameters that match the parental system (temperature: 26°C; pH: 8.1-8.2; salinity: 35 parts-per-thousand), dosed with methylene blue (500 µL) and heated by a glass aquarium heater. Eggs are still attached to pot shards and propped upright in zebrafish removable tank inserts within the larger aquarium.

While incubating *A. ocellaris* eggs, it is essential to provide adequate water flow to maintain aeration and prevent debris from settling on eggs. In nature, this care is conveyed by the fanning and mouthing of eggs by parents and can be easily imitated using a wooden air diffuser (connected via hose to an air pump) placed in front of the eggs (see below image for setup). By fitting an attenuator/dial on the air hose the airflow can be easily adjusted to an optimal level, where eggs are gently waving but not violently shaking. It is important to manually remove any deceased eggs from the pot shards that inevitably lead to reduced egg-movement, spread of disease, and increased mortality. By 4 dpf mortality rates usually stabilise and reduce the need for manual cleaning.

NOTE: returning any (injected or non-injected) eggs to parents will most likely result in their rejection and/or consumption by the parents. Although we have had no success in returning eggs to any of our parental pairs, this may vary across individual pairs and be worth trialling on a case-by-case basis.

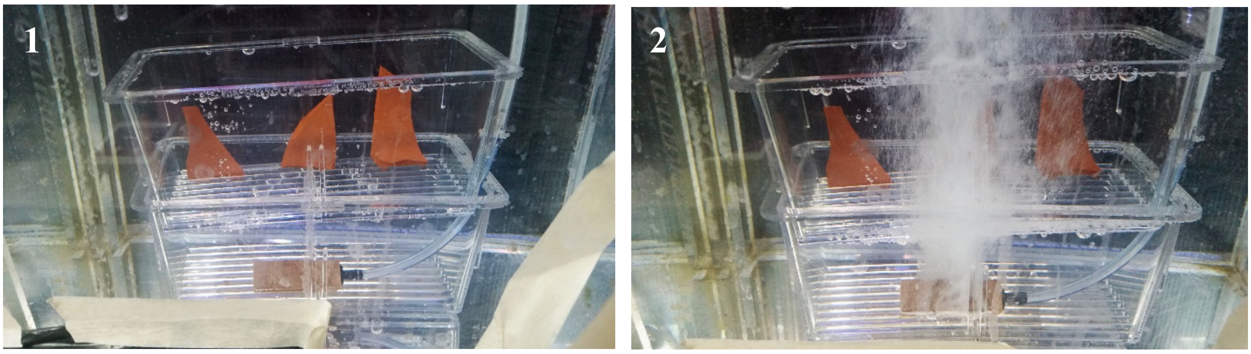


Image 1: incubation setup immediately following injections, with shards containing eggs positioned upright in a standard zebrafish removable tank insert. A wooden air diffuser was positioned underneath and ~10 cm in front of the shards.  Image 2: throughout incubation eggs were kept aerated and gently swaying using moderate flow from the fine air bubbles produced by a wooden air diffuser.

7-8 dpf, the silvering of embryos signal they are nearing ready-to-hatch that evening. It is during this period when a few environmental changes are essential to induce hatching that includes *1)* moving the eggs to a new tank with clean water and eliminated light by lining the sides and top of the aquarium with black-wrap or plastic, *2)* greater egg disturbance via increased airflow and/or placing airflow closer to the eggs, and *3)* the introduction of algal paste to create green-water and rotifer culture as a food source for larvae. Most eggs should hatch over the next couple of nights.

Post-hatch, the airflow must be reduced to a small stream to provide adequate aeration whilst not harming the fragile newly hatched larvae. Any non-hatched eggs should be removed at this time. During this time, it is safe to introduce diffused down-welling light using a thin towel over the aquarium lid; however, the sides of the aquarium should remain fully blacked out. An optimal rotifer density of about 8-10 $mL^{-1}$ should be maintained for 7 days. Depending on larval density and hatching success, the rotifer population may co-culture and not require any further addition, if green water is maintained by the dosing of algal paste. After the fifth day, newly (24 h) hatched *artemia* nauplii can be introduced as a food to provide greater nourishment. Beyond the seventh day post-hatch (dph), all larvae should be capable of feeding exclusively on *artemia* nauplii. After phasing out rotifers from the diet, it is no longer necessary to add green water. Throughout the rearing process, it is crucial to ensure the ammonia level does not rise to become too toxic (≤0.2 ppm, preferably undetectable), and is best kept low by having a constant drip feed of new salt water. Any dead larvae and/or debris should be carefully siphoned out.

Upon reaching the metamorphosis stage (10-12 dph), the larvae will have formed stripes and settled. At this stage, a few system improvements can be safely made including the installation of an air-driven sponge filter to provide filtration, and objects for shelter (e.g. plastic pipes, small pots, anemones). The diet of settled larvae and early-juveniles can be shifted towards fine-sized commercial fish pellets or flakes. Adult-sized pellets can be used but must be crushed into finer pieces for young fish to swallow.

### 2. sgRNA sequences

*RH2B* sgRNA

Target sequence 1: *RH2B* Exon 4 (‘5 – ‘3) GGCCAUGGAGACAGCUGAGA

Target sequence 2: *RH2B* Exon 5 (‘5 – ‘3) UGGAAUGGGUGGCAUGGUUG

Target sequence 3: *RH2B* Exon 5 (‘5 – ‘3) CUGAGCACCAUUGGAAUGGG

<u>*tyr* sgRNA</u>

Target sequence 1: *tyr* Exon 2 ('5 – '3) GGCGUUGUGUAACGCAACA

Target sequence 2: *tyr* Exon 2 ('5 – '3) GAGGAGUACAACAGCCGAG

**3. sgRNA *in-vitro* assay**

For further details on the *in-vitro* assay (e.g. reaction buffer ingredients) see the original protocol outlined by Grainger *et al.* 2017.

Reaction mix:

| | |
|---|---|
| PCR product | 8 µL |
| 10X Cas9 reaction buffer | 2 µL |
| Cas9 protein | 2 µL |
| sgRNA | 1 µg |
| Nuclease free water to | 20 µL |

Incubation steps:

1. Incubate the above mix at 37°C for 1 h, and 98°C for 20 min to denature excess Cas9 protein. Store at -4°C.
2. Run on a 2.5-3.0% agarose gel to analyse cleavage activity Cas9 directed by your sgRNA by comparison with PCR product without either Cas9 or sgRNA.

**4. Crafting of microneedles**

Microneedles for CRISPR-construct delivery were pulled from borosilicate capillaries (Harvard Apparatus: 1.0x0.58x100 mm) using a micropipette puller (P-1000 Sutter Instruments with a 2 mm wide trough filament) to produce microneedles with a short (3-4 mm) taper. This was found to be important in minimising the bending of the needle tip when attempting to penetrate the thick chorion of anemonefish eggs. Our micropipette puller settings were as follows: Heat: +10% ramp value, Pull: 55, Vel.: 70, Time: 165, Pressure: 500.

Microneedles were viewed under a microscope against a 0.01mm scale calibration slide, and tips were broken back using a pair of fine-tipped forceps to give a final tip diameter between 10-15 μm.

**5. Calibration of microinjector settings**

Pre-cut needles were backloaded with 2 μL of phenol red dye mixed with water and fastened on the microinjector (Narishige IM-400). Air pressure and pulse duration settings were adjusted to produce a single droplet with a diameter of 125-140 μm measured on a calibration slide (0.01 mm scale) that roughly corresponded to a volume between 1-1.4 nL. Our used injector settings were as follows: Pressure: 1.017 psi, Time: 0.8-1.0 sec., Back pressure: 0 psi (on).

**6. PCR primer sequences**

*RH2B*

| | |
|---|---|
| Forward primer ('5 – '3) | TCCCAGTACACAACGCAGTC |
| Reverse primer ('5 – '3) | AGGGTGTCTGGAGATGTGGA |

*tyr*

| | |
|---|---|
| Forward primer ('5 – '3) | ACCAGTCGACACTTGCTGCT |
| Reverse primer ('5 – '3) | TCGGTCCATAGGTCCCGTCT |

### 7. PCR routine

See New England Biolabs (product code: M0496S) for recommended reagent quantities for a desired reaction volume.

Thermocycler steps:

| | |
|---|---|
| Initial denaturation | 95°C (for 30 seconds) |
| Repeated for 30 cycles | 95°C (15 seconds) |
| | 60°C (60 seconds) |
| | 68°C (30 seconds) |
| Final extension | 68°C (5 minutes for *in-vitro* test; 10 minutes for subcloning) |
| Hold | 4°C (forever) |

### 8. Frameshift mutation analysis

| WT *RH2B* AA no. | 234 | 235 | 236 | 237 | 238 | 239 | 240 | 241 | 242 | 243 | 244 | 245 | 246 | 247 | 248 | 249 | 250 | 251 | 252 | 253 | 254 | 255 | 256 | 257 | 258 |
|---|---|---|---|---|---|---|---|---|---|---|---|---|---|---|---|---|---|---|---|---|---|---|---|---|---|
| WT | A | A | A | Q | Q | Q | D | S | A | S | T | Q | K | A | E | K | E | V | T | R | M | C | V | L | M |
| M1.1 | A | A | A | Q | Q | Q | D | S | A | S | T | Q | K | A | E | K | E | V | T | R | M | C | V | L | M |
| M1.2 | A | A | A | Q | Q | Q | D | S | A | S | T | Q | K | A | E | K | E | V | T | R | M | C | V | L | M |
| M4.1 | A | A | A | Q | Q | Q | D | S | A | S | T | Q | K | A | E | K | E | V | T | R | M | C | V | L | M |
| M4.2 | A | A | A | Q | Q | Q | D | S | A | S | T | Q | K | A | E | K | E | V | T | R | M | C | V | L | M |
| M4.3 | A | A | A | Q | Q | Q | D | S | A | S | T | Q | K | A | E | K | E | V | T | R | M | C | V | L | M |
| Bovine *RH1* | A | A | A | Q | Q | Q | E | S | A | T | T | Q | K | A | E | K | E | V | T | R | M | V | I | I | M |
| Bovine *RH1* AA no. | 233 | 234 | 235 | 236 | 237 | 238 | 239 | 240 | 241 | 242 | 243 | 244 | 245 | 246 | 247 | 248 | 249 | 250 | 251 | 252 | 253 | 254 | 255 | 256 | 257 |

| WT *RH2B* AA no. | 259 | 260 | 261 | 262 | 263 | 264 | 265 | 266 | 267 | 268 | 269 | 270 | 271 | 272 | 273 | 274 | 275 | 276 | 277 | 278 | 279 | 280 | 281 | 282 | 283 |
|---|---|---|---|---|---|---|---|---|---|---|---|---|---|---|---|---|---|---|---|---|---|---|---|---|---|
| WT | V | F | G | F | L | F | A | W | T | P | Y | A | S | F | A | A | W | I | F | F | N | K | G | A | A |
| M1.1 | V | F | G | F | L | F | A | W | T | P | Y | A | S | F | A | A | W | I | F | F | N | K | G | A | A |
| M1.2 | V | F | G | F | L | F | A | W | T | P | Y | A | S | F | A | A | W | I | F | F | N | K | G | A | A |
| M4.1 | V | F | G | F | L | F | A | W | T | P | Y | A | S | F | A | A | W | I | F | F | N | K | G | A | A |
| M4.2 | V | F | G | F | L | F | A | W | T | P | Y | A | S | F | A | A | W | I | F | F | N | K | G | A | A |
| M4.3 | V | F | G | F | L | F | A | W | T | P | Y | A | S | F | A | A | W | I | F | F | N | K | G | A | A |
| Bovine *RH1* | V | I | A | F | L | I | C | W | L | P | Y | A | G | V | A | F | Y | I | F | T | H | Q | G | S | D |
| Bovine *RH1* AA no. | 258 | 259 | 260 | 261 | 262 | 263 | 264 | 265 | 266 | 267 | 268 | 269 | 270 | 271 | 272 | 273 | 274 | 275 | 276 | 277 | 278 | 279 | 280 | 281 | 282 |

| WT *RH2B* AA no. | 284 | 285 | 286 | 287 | 288 | 289 | 290 | 291 | 292 | 293 | 294 | 295 | 296 | 297 | 298 | 299 | 300 | 301 | 302 | 303 | 304 | 305 | 306 | 307 | 308 |
|---|---|---|---|---|---|---|---|---|---|---|---|---|---|---|---|---|---|---|---|---|---|---|---|---|---|
| WT | F | S | A | V | S | M | A | I | P | A | F | F | S | K | S | S | A | L | F | N | P | V | I | Y | I |
| M1.1 | F | - | C | L | H | G | H | P | C | F | L | L | K | E | F | S | F | V | Q | S | C | Y | L | H | P |
| M1.2 | F | S | C | L | H | G | H | P | C | F | L | L | K | E | F | S | F | V | Q | S | C | Y | L | H | P |
| M4.1 | - | S | C | L | H | G | H | P | C | F | L | L | K | E | F | S | F | V | Q | S | C | Y | L | H | P |
| M4.2 | F | S | C | L | H | G | H | P | C | F | L | L | K | E | F | S | F | V | Q | S | C | Y | L | H | P |
| M4.3 | F | F | - | - | - | - | - | - | - | - | - | - | - | - | - | - | R | C | S | I | L | L | S | T | S |
| Bovine *RH1* | F | G | P | I | F | M | T | I | P | A | F | F | A | K | T | S | A | V | Y | N | P | V | I | Y | I |
| Bovine *RH1* AA no. | 283 | 284 | 285 | 286 | 287 | 288 | 289 | 290 | 291 | 292 | 293 | 294 | 295 | 296 | 297 | 298 | 299 | 300 | 301 | 302 | 303 | 304 | 305 | 306 | 307 |

| WT *RH2B* AA no. | 309 | 310 | 311 | 312 | 313 | 314 | 315 | 316 | 317 | 318 | 319 | 320 | 321 | 322 | - | - | 323 | 324 | 325 | 326 | 327 | 328 | 329 | 330 | 331 |
|---|---|---|---|---|---|---|---|---|---|---|---|---|---|---|---|---|---|---|---|---|---|---|---|---|---|
| WT | L | M | N | K | Q | F | R | N | C | M | L | S | T | I | - | - | G | M | G | G | M | V | E | D | E |
| M1.1 | Y | E | Q | T | V | P | * | L | H | A | E | H | H | W | N | G | W | H | G | * | G | * | D | L | S |
| M1.2 | Y | E | Q | T | V | P | * | L | H | A | E | H | H | W | N | G | W | H | G | * | G | * | D | L | S |
| M4.1 | Y | E | Q | T | V | P | * | L | H | A | E | H | H | W | N | G | W | H | G | * | G | * | D | L | S |
| M4.2 | Y | E | Q | T | V | P | * | L | H | A | E | H | H | W | N | G | W | H | G | * | G | * | D | L | S |
| M4.3 | L | * | T | N | S | S | V | T | A | C | * | A | P | L | E | W | V | A | W | L | R | M | R | P | Q |
| Bovine *RH1* | M | M | N | K | Q | F | R | N | C | M | V | T | T | L | C | C | G | K | N | P | L | G | D | D | E |
| Bovine *RH1* AA no. | 308 | 309 | 310 | 311 | 312 | 313 | 314 | 315 | 316 | 317 | 318 | 319 | 320 | 321 | 322 | 323 | 324 | 325 | 326 | 327 | 328 | 329 | 330 | 331 | 332 |

| WT *RH2B* AA no. | 332 | 333 | 334 | 335 | 336 | 337 | 338 | 339 | 340 | 341 | 342 | 343 | 344 | 345 | - | - | 346 |
|---|---|---|---|---|---|---|---|---|---|---|---|---|---|---|---|---|---|
| WT | T | S | V | S | T | S | K | T | E | V | S | S | V | S | - | - | * |
| M1.1 | I | Y | Q | Q | D | R | S | - | L | L | S | V | L | - | - | - | - |
| M1.2 | I | Y | Q | Q | D | R | S | - | L | L | S | V | L | - | - | - | - |
| M4.1 | I | Y | Q | Q | D | R | S | - | L | L | S | V | L | - | - | - | - |
| M4.2 | I | Y | Q | Q | D | R | S | - | L | L | S | V | L | - | - | - | - |
| M4.3 | Y | L | P | A | R | Q | K | S | P | Q | C | P | - | - | - | - | - |
| Bovine *RH1* | A | S | T | T | V | S | K | T | E | T | S | Q | V | A | P | A | * |
| Bovine *RH1* AA no. | 333 | 334 | 335 | 336 | 337 | 338 | 339 | 340 | 341 | 342 | 343 | 344 | 345 | 346 | 347 | 348 | 349 |

*Supplementary Figure. 1.* Translated sequence alignment of frameshifted alleles found in *RH2B*-M1 and -M4, and wildtype (WT) *RH2B* for reference. Sequences were aligned against bovine rhodopsin (*RH1*), as an opsin template. The chromophore binding site (bovine *RH1* AA no., Lys296) is boxed in blue. Amino acid (AA) numbering schemes were according to WT *RH2B* (upper) and bovine *RH1* (lower). Translated sequences were aligned using MAFFT Alignment (*v7.450*) in Geneious.

Katoh, K., & Standley, D. M. (2013). MAFFT Multiple Sequence Alignment Software Version 7: Improvements in Performance and Usability. *Molecular Biology and Evolution*, *30*(4), 772–780. https://doi.org/10.1093/molbev/mst010
